# AMPK activation restores salivary function following radiation treatment

**DOI:** 10.1101/2022.11.29.518381

**Authors:** Rachel Meyer, Kristy Gilman, Brenna A. Rheinheimer, Lauren Meeks, Kirsten H. Limesand

## Abstract

Head and neck cancers represent a significant portion of cancer diagnoses, with head and neck cancer incidence increasing in some parts of the world. Typical treatment of early-stage head and neck cancers includes either surgery or radiotherapy; however, advanced cases often require surgery followed by radiation and chemotherapy. Salivary gland damage following radiotherapy leads to severe and chronic hypofunction with decreased salivary output, xerostomia, impaired ability to chew and swallow, increased risk of developing oral mucositis, and malnutrition. There is currently no standard of care for radiation-induced salivary gland dysfunction and treatment is often limited to palliative treatment that provides only temporary relief. AMP-activated protein kinase (AMPK) is an enzyme that activates catabolic processes and has been shown to influence the cell cycle, proliferation, and autophagy. In the present study, we found that radiation (IR) treatment decreases tissue levels of phosphorylated AMPK following radiation and decreases intracellular NAD+ and AMP while increasing intracellular ATP. Further, expression of Sirtuin-1 (SIRT1) and nicotinamide phosphoribosyl transferase (NAMPT) were lower five days following IR. Treatment with AMPK activators, AICAR and metformin, attenuated compensatory proliferation (days 6, 7 and 30) following IR, and reversed chronic (day 30) salivary gland dysfunction post-IR. Additionally, treatment with metformin or AICAR increased markers of apical/basolateral polarity (phosphorylated aPKCζT560 positive area) and differentiation (amylase positive area) within irradiated parotid glands to levels similar to untreated controls. Taken together, these data suggest that AMPK may be a novel therapeutic target for treatment of radiation-induced salivary damage.

## Introduction

Approximately 55,000 new cases of head and neck cancer will be diagnosed next year in the United States (Siegel et al. 2022). Despite increasing incidence of head and neck cancers, treatment options remain limited. Standard treatment often includes surgery and radiotherapy (Cramer et al. 2019). Although treatment options and survival rates vary depending on location and severity of malignancy, thousands of cancer survivors retain impaired quality of life following treatment due to secondary damage to surrounding tissues, including salivary glands. Radiotherapy-induced salivary gland damage chronically impacts function, typically resulting in decreased salivary output, difficulty eating, oral mucositis, and malnutrition (Grundmann et al. 2009). Treatment options are commonly reserved to palliative care, with few options available to directly treat or prevent damage to the gland (Jensen et al. 2019).

Compensatory proliferation in the wound healing response is a process conserved across tissue types and has been implicated in the cellular response to radiation. While proliferation following tissue damage is typically considered beneficial to replace apoptotic and/or necrotic cells, the proliferative response in radiation-treated salivary glands occurs concomitantly with significant reductions in salivary output (Grundmann et al. 2010; Morgan-Bathke et al. 2014). In addition to compensatory proliferation, cytoskeletal disruptions and disorganization of the Par3-aPKC polarity complex have been associated with deficient wound healing and correlates with a significant decrease in saliva production (Abreu-Blanco et al. 2012; Chibly et al. 2018; Niessen et al. 2013; Wong et al. 2018). Previous work indicated that administration of insulin-like growth factor-1 (IGF-1) to mice following exposure to 5-Gy radiation restores glandular function and salivary output (Chibly et al. 2018; Grundmann et al. 2010;). Functional restoration of the gland is associated with attenuation of proliferative responses in IGF-1 treated irradiated mice (Grundmann et al. 2010), possibly indicating that the newly dividing cells have limited functional capability and are therefore not contributing to saliva production. Taken together, these data suggest that reducing proliferative cells in the parotid salivary gland may play a role in improving functionality following radiation therapy.

One of the most frequently studied metabolic regulators, AMP-activated protein kinase (AMPK), modulates flux through metabolic pathways in response to low cellular energy states (Hardie et al. 1998; Ke et al. 2018). AMPK is activated in a two-step process involving increased binding of AMP and phosphorylation on threonine^172^ by LKB1 or CAMKK2 (Herzig and Shaw 2018). Active AMPK increases production of NAD+ via inducing expression of nicotinamide phosphoribosyltransferase (NAMPT), a key enzyme in NAD biosynthesis, (Garten et al. 2015); the AMPK-mediated increase in NAD+ enhances the activity of the NAD-dependent histone deacetylase sirtuin-1 (SIRT1) (Cantó et al. 2009) that plays a regulatory role in autophagy induction (Lee 2019). Previous work has implicated autophagy in amelioration of radiation-induced salivary gland dysfunction (Morgan-Bathke et al. 2015). Autophagy deficient mice have a more severe response to targeted head and neck radiation, with decreased acute and chronic salivary flow rates following 5-Gy radiation. While autophagy is not acutely activated following IR, IGF-1 pretreatment induces autophagy and improves salivary function. Autophagy induction likely acts, in part, via inhibition of mTOR signaling, which is increased 4-5 days following IR. Indeed, irradiated mice treated with the rapalogue CCI-779 have improved salivary function 30 days following radiation; parotid salivary glands from these mice also show fewer proliferative cells compared to irradiated controls (Morgan-Bathke et al. 2014). AMPK activation is also known to activate autophagy, in part, due to inhibition of mammalian target of rapamycin (mTOR1) signaling (Imamura et al. 2001; Jones et al. 2005). While AMPK activation has not been studied in the salivary gland following radiation exposure, treatment with metformin, a drug known to activate AMPK, improves salivary flow rate in a mouse model of Sjögren’s syndrome (Kim et al. 2019), indicating a potential therapeutic use for metformin in the salivary gland response to radiation. Taken together, the role of AMPK in the inhibition of proliferation and activation of autophagy implicates this enzyme as a prime molecular target to improve radiation-induced salivary gland dysfunction.

Based on preliminary evidence suggesting a role for AMPK in salivary secretion, the present study aimed to 1) asses changes in NAD+ and AMP levels in parotid salivary glands following IR, 2) characterize the levels of phosphorylated AMPK and enzymes in downstream pathways following IR, 3) determine the effect of pharmacological activation of AMPK with AICAR or metformin on radiation-induced compensatory proliferation in parotid cells and 4) evaluate the effect of AICAR or metformin treatment on chronic salivary output following radiation exposure. We hypothesize that treatment with AICAR or metformin will attenuate the well-characterized proliferative response following IR and pharmacological AMPK activation with AICAR or metformin will improve salivary output when administered following radiation.

## Materials and Methods

### Mice

FVB mice were purchased from Jackson Laboratories (Bar Harbor, ME) and maintained in accordance with protocols approved by the University of Arizona Institutional Animal Care and Use Committee. This study conforms to the ARRIVE guidelines. Administration of metformin (Sigma-Aldrich, St. Louis, MO) or AICAR (Toronto Research Chemicals, North York, Ontario, Canada) was performed via oral gavage (metformin, 100 mg/kg body weight) or intraperitoneal injection (AICAR, 500 mg/kg body weight).

### Radiation treatment

The head and neck region of mice received single (5-Gy) or fractionated (2Gy/day X5 days) dose of radiation using a ^60^Cobalt Teletherapy instrument (Atomic Energy of Canada Limited Theratron-80) as previously described (Chibly et al. 2018)(Limesand et al. 2010).

NAD+ and AMP Metabolite Analysis. Mice were unirradiated or received a 5-Gy dose of radiation. Parotid glands removed at day 5 following IR, snap frozen, and sent to Metabolon, Inc (Morrisville, NC) for metabolomics analysis by LC-MS as previously described (Meeks et al. 2021).

ATP cell assay. ATP concentration was determined by ATP bioluminescence assay (Roche, Tucson, AZ) following the manufacturers’ instructions. See appendix for more information.

AMP and ATP tissue assay. Parotid glands were removed, one gland was homogenized in 400ul AMP assay buffer while the second gland was homogenized in 100ul ATP assay buffer (Abcam, Cambridge, United Kingdom). AMP and ATP concentration were determined following the manufacturers’ instructions.

### Western Blot

Parotid glands were removed, protein extracted, and SDS-PAGE conducted as previously described (Gilman et al. 2021). Refer to the Appendix for antibody information.

### RNA isolation and qRT-PCR

Parotid glands were removed from mice at days 3, 5 and 30 after IR. RNA was isolated as previously described (Gilman et al. 2019). Refer to the Appendix for primer information.

### Blood Glucose

Following a 4-hour fast, the distal 1 mm of tail was cut, and glucose was measured from 10-20 µL whole blood by CVS Health Advanced Blood Glucose Meter (CVS Health, Woonsocket, RI).

Histology and Immunofluorescent staining. Salivary glands were removed, fixed in 10% (v/v) formalin for 24 hours and sent to IDEXX Bioresearch (Columbia, MO) for embedding. Immunofluorescent staining and quantification were performed as described in the Appendix.

### Saliva collection

Stimulated whole saliva collection was performed on days 3 and 30 following IR as previously described (Gilman et al. 2021).

### Statistical analysis

Statistical analysis was performed using GraphPad Prism 6 software (GraphPad Software, San Diego, CA). Manual cell counts from immunofluorescent stained slides were analyzed by ANOVA with Tukey’s multiple comparisons test. Percent positive staining area was analyzed by ANOVA followed by Bonferroni’s *post hoc* comparisons. qRT-PCR data was normalized to loading controls (15S) and analyzed with a t-test comparing irradiated values to untreated. Densitometry data from Western blot was analyzed using a t-test. Saliva collection data was normalized to untreated values for each collection day and compared with a one-way ANOVA with Newman-Keuls multiple comparisons test. Treatment groups denoted with different letters above the graph are significantly different from each other while data from groups without statistical differences are indicated with the same letter.

## Results

### Radiation treatment leads to suppression of the AMPK pathway

AMPK is primarily activated in response to changes in intracellular metabolism, specifically an increase in AMP:ATP ratio and decreased NAD+ availability. Radiation exposure has been shown to affect metabolism within a whole organism or specific tissue (Golla et al. 2017; Manna et al. 2013). To determine the effect of radiation on metabolites involved in AMPK activation, untargeted metabolomics was performed on parotid glands from irradiated and untreated (UT) mice when compensatory proliferation begins (day 5 following IR) (Meeks et al. 2021). We observed a ∼50% decrease in NAD+ levels and a ∼30% decrease in AMP levels in irradiated parotid glands compared to untreated controls (Fig. 1A,B). Further, radiation significantly increased ATP levels in parotid glands (Fig. 1C). To determine the effect of radiation on AMPK activation, we measured phosphorylated (activated) AMPK^T172^ at days 3 and 5 following IR. Phosphorylated-AMPK protein levels decreased 3 and 5 days following IR in parotid glands compared to untreated (Fig. 1D,E). AMPK activity has also been shown to regulate nicotinamide phosphoribosyltransferase (NAMPT) expression and Sirtuin-1 (SIRT1) activity (Brandauer et al. 2013; Cantó et al. 2009). SIRT1 and NAMPT expression are significantly lower in mouse parotid salivary glands at day 5 following radiation (Fig. 1F). These data suggest radiation impacts cellular energy homeostasis at early time points leading to a reduction in AMPK signaling.

**Figure 1.**
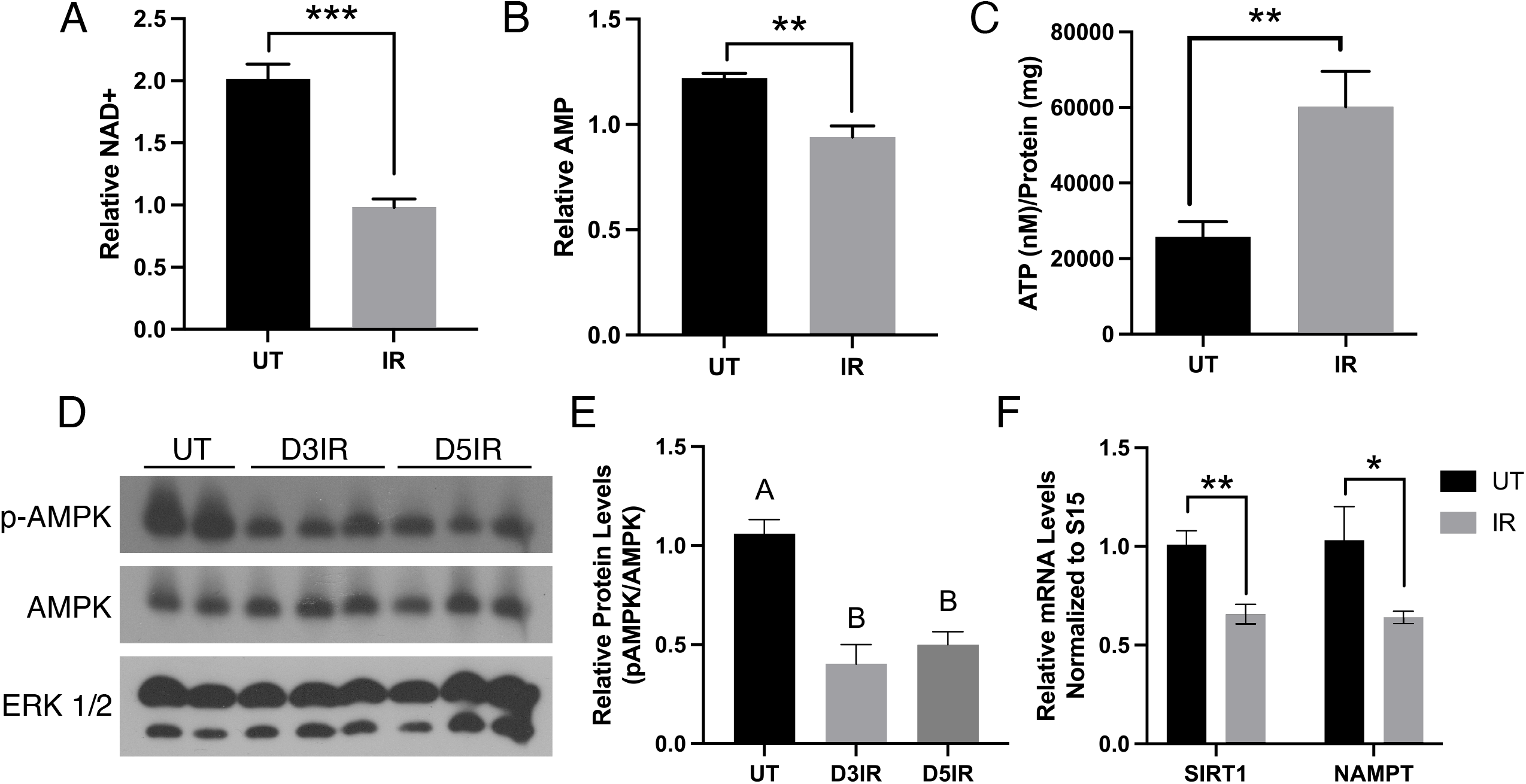
Radiation treatment leads to suppression of the AMPK pathway. FVB mice were untreated (UT) or exposed to 5-Gy radiation (IR) and sacrificed at days 3 or 5 following radiation. (A) NAD+ and (B) AMP levels were determined via LC-MS in untreated (n=4) or irradiated (n=4) parotid salivary glands. (C) Primary cultures of parotid gland cells were prepared and left untreated (n=6) or received 5-Gy IR (n=5). Cell lysates were collected at day 3 post-IR and intracellular ATP levels were determined with a luciferin-luciferase assay. (D) Protein was extracted from untreated and day 3 or 5 irradiated parotid glands and levels of phosphorylated AMPK (p-AMPK), total AMPK (t-AMPK) and ERK 1/2 (loading control) were determined by immunoblot. (E) Densitometry data from untreated (n=4), day 3 (n=3) or day 5 (n=3) irradiated parotid glands. AMPK (p-AMPK) levels were normalized to total AMPK (t-AMPK) levels and are presented as mean ± SEM. Statistical differences were determined with a one-way ANOVA followed by Tukey’s post-hoc test. Groups with different letter designations are significantly different from each other, p<0.05. (F) cDNA was prepared from untreated (n=4) or 5-Gy irradiated parotid glands at day 5 following radiation (n=4) and qRT-PCR was performed with primers specific to sirtuin-1 (SIRT1) and nicotinamide phosphoribosyltransferase (NAMPT). Data was normalized to 15S ribosomal RNA as an internal control and calculated as fold-change relative to the average of untreated mice. (A-C, F) Data are presented as mean ± SEM and statistical differences between groups were determined with an unpaired t-test, *p<0.05, **p<0.01.

### Treatment with metformin or AICAR following radiation activates AMPK in parotid glands

To confirm the AMPK-activating effect of AICAR and metformin in our model of radiation, we treated irradiated mice with AICAR or metformin at days 4, 5, and 6 following radiation. We measured intracellular AMP and ATP levels at days 7 and 30 following radiation and found no significant differences in AMP levels between groups (Fig. 2 A,B; Supplemental Fig. 1A,B). Mice treated with AICAR or metformin had lower fasting blood glucose at day 7 following IR compared to UT (Fig. 2C), with no differences between groups at day 30 (Supplemental Fig. 1C). To confirm that AICAR and metformin treatment activate AMPK in parotid glands, we measured phosphorylated AMPK at days 7 and 30 following IR. Radiation decreases phospho-AMPK in parotid glands at day 7 compared to UT, while AICAR or metformin treatment increase phospho-AMPK levels compared to IR (Fig. 2D,E). At day 30, phospho-AMPK levels are similar between UT and IR; however, metformin or AICAR treatment significantly increase phospho-AMPK levels compared to IR (Supplemental Fig. 1D,E). There were no differences between groups in the expression of SIRT1 and NAMPT at day 30 following radiation (Supplemental Fig. 1F,G). These data indicate that administration of AICAR or metformin following radiation induces AMPK activation independent of changes to intracellular AMP and ATP levels.

**Figure 2.**
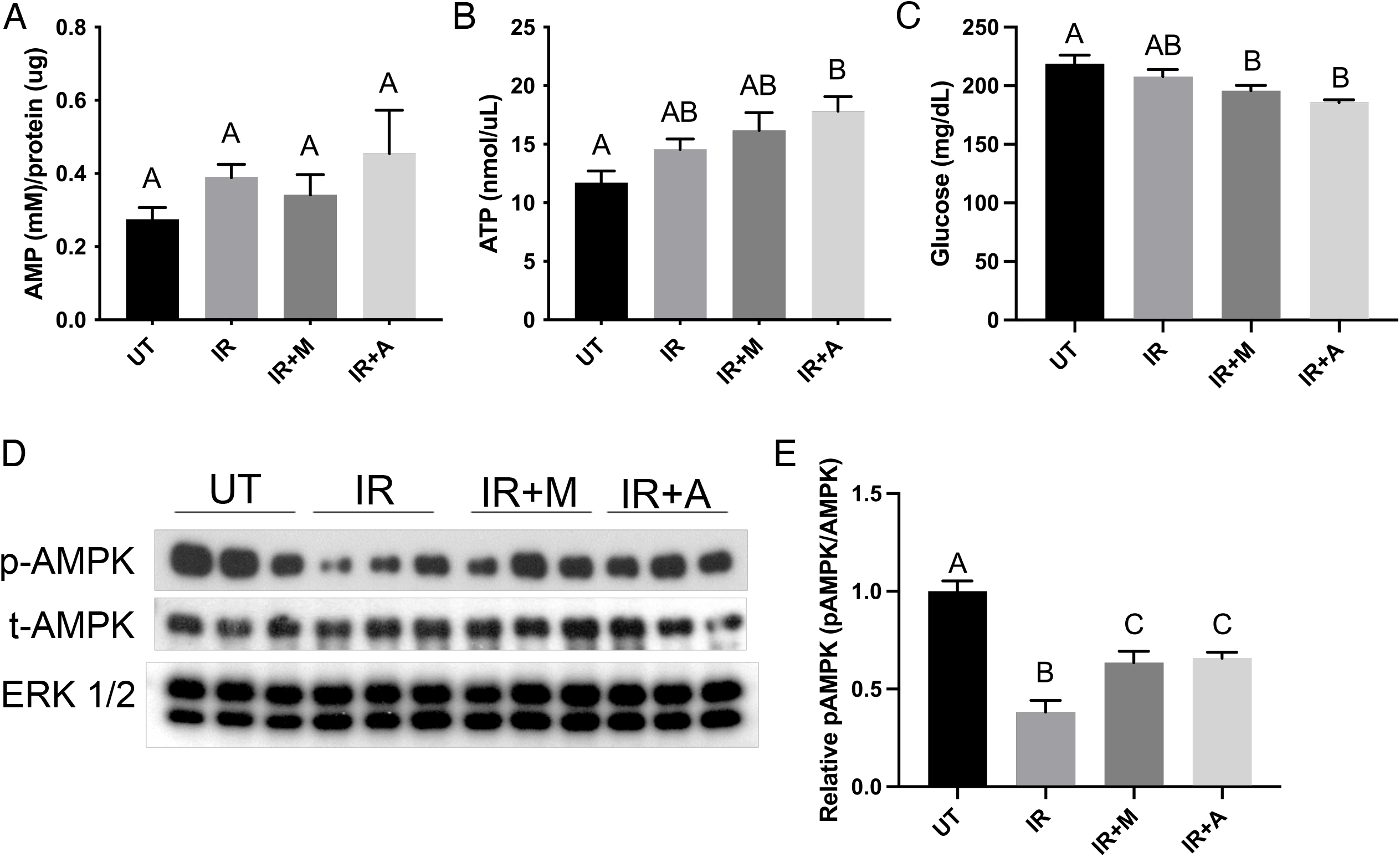
Treatment with metformin or AICAR following radiation activates AMPK in parotid glands. FVB mice were untreated (UT, n=4) or exposed to 5-Gy radiation (n=4) with additional mice receiving AICAR (n=3) or metformin (n=4) injections on days 4, 5 and 6 after radiation. Parotid salivary glands were collected at day 7 following radiation. Levels of (A) AMP and (B) ATP were evaluated from tissue lysates via colorimetric assays as described in Materials and Methods. (C) Blood was collected from mice in indicated treatment groups on day 7 post-IR and glucose levels were measured via handheld glucometer following a 4-hour fast. (D-E) Levels of phosphorylated AMPK (p-AMPK), total AMPK (t-AMPK) and ERK 1/2 (loading control) were evaluated via immunoblot. AMPK (p-AMPK) levels were normalized to total AMPK (t-AMPK) levels. (A-E) Data are presented as mean ± SEM. Statistical differences were determined with a one-way ANOVA followed by Tukey’s post-hoc test. Groups with different letter designations are significantly different from each other, p<0.05.

### Treatment with AICAR or metformin decreases compensatory proliferation following radiation

Previous research restoring salivary gland function demonstrated a decrease in compensatory proliferation following IR that occurs concomitantly with improved salivary output (Grundmann et al. 2010; Morgan-Bathke et al. 2015). Further, AMPK activation inhibits proliferation by decreasing flux through anabolic pathways that require excessive ATP, inhibition of mTOR signaling and activation of p53 (Agarwal et al. 2015; Hao et al. 2017; Jones et al. 2005). To determine if AMPK activation has a similar effect on proliferation in the parotid salivary gland, we treated mice with one dose of AICAR or metformin 5 days following IR and collected glands 24 hours later (day 6 after IR) for Ki67 staining. Metformin and AICAR treatment attenuated the proliferative response to radiation, exhibiting a 65% and 63% decrease in Ki67+ cells compared to IR alone, respectively (Fig. 3A-E). Given that previous models of functional restoration have administered multiple doses of therapeutic compound to reverse salivary gland dysfunction, mice were treated with 3 doses of AICAR or metformin at days 4, 5, and 6 following radiation and glands were collected 24 hours after the last dose (day 7 after IR; Fig. 3F-J). Treatment with metformin or AICAR over 3 days decreases the proliferative response to radiation at day 7 following IR by 43% and 40%, respectively (Fig. 3F-J). These data indicate that AMPK activation following IR attenuates the proliferative response to radiation in parotid salivary glands.

**Figure 3.**
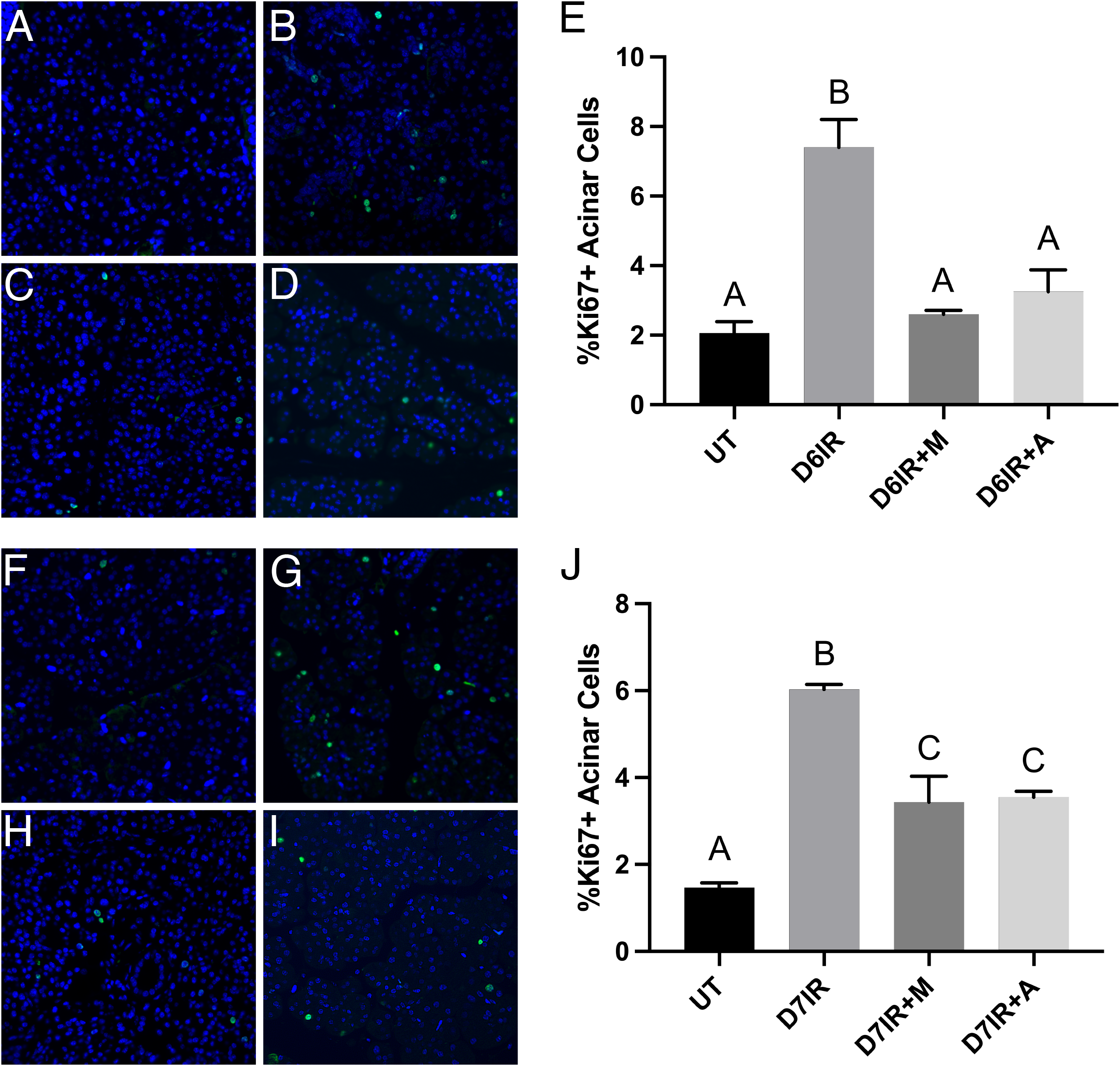
AICAR or metformin treatment decreases compensatory proliferation of parotid glands following radiation. FVB mice were untreated or exposed to 5-Gy radiation with a subset of mice receiving AICAR (500mg/kg) or metformin (100 mg/kg) on day 5 (A-E) or days 4, 5 and 6 (F-J). Parotid salivary glands were collected at indicated timepoints and immunohistochemistry for Ki67 was performed. (A-D) Representative images of Ki67 positive cells (green) over total nuclei stained with DAPI (blue) in (A) untreated (n=4), (B) day 6 following 5-Gy radiation (n=4), (C) day 6 following 5-Gy radiation and metformin on day 5 (100 mg/kg, 1 dose), or (D) day 6 following 5-Gy radiation and AICAR on day 5 (500 mg/kg, 1 dose) (n=5). (E) Quantification of Ki67+ nuclei as a percentage of the total number of nuclei from 5 fields of view per mouse. (F-I) Representative images of Ki67 positive cells (green) over total nuclei stained with DAPI (blue) in (F) untreated (n=4), (G) day 7 following 5-Gy radiation (n=4), (H) day 7 following 5-Gy radiation and metformin on days 4, 5, and 6 (100 mg/kg, 3 doses) (n=4), or (I) day 7 following 5-Gy radiation and AICAR on days 4, 5, and 6 (500 mg/kg, 3 doses) (n=4). (J) Data are presented as mean ± SEM and statistical differences were determined with a one-way ANOVA followed by Tukey’s post-hoc test, p<0.05.

### AICAR or metformin treatment increases salivary output, increases amylase and phosphorylated aPKCζ area and decreases proliferation 30 days post-treatment

Reduced proliferation in irradiated salivary glands correlates with improved salivary output in previous models of restoration (Chibly et al. 2018; Grundmann et al. 2010; Morgan-Bathke et al. 2015). To determine the restorative potential of pharmacological AMPK activation, we irradiated mice and treated with AICAR or metformin (Fig. 4A). Radiation treatment decreases salivary output by 29% or 26% at days 3 or 30, respectively, following IR compared to UT (Fig. 4B,C). Treatment with metformin or AICAR improves salivary output compared to IR, with no differences from UT control salivary output (Fig. 4C). Non-irradiated mice treated with three doses of AICAR have no difference in salivary flow compared to UT; however, non-irradiated mice treated with metformin have lower salivary flow rates compared to UT (Supplemental Fig. 2). To determine the effect of metformin or AICAR treatment on salivary output with fractionated doses of radiation, we treated mice with 2Gy/day for 5 days and administered metformin or AICAR at days 4, 5, and 6 following the final radiation dose (Fig. 4D). Fractionated radiation decreases salivary output at day 30 following radiation by 33% compared to UT, while metformin or AICAR treatment improves salivary output at day 30, with no significant differences from UT (Fig. 4E).

**Figure 4.**
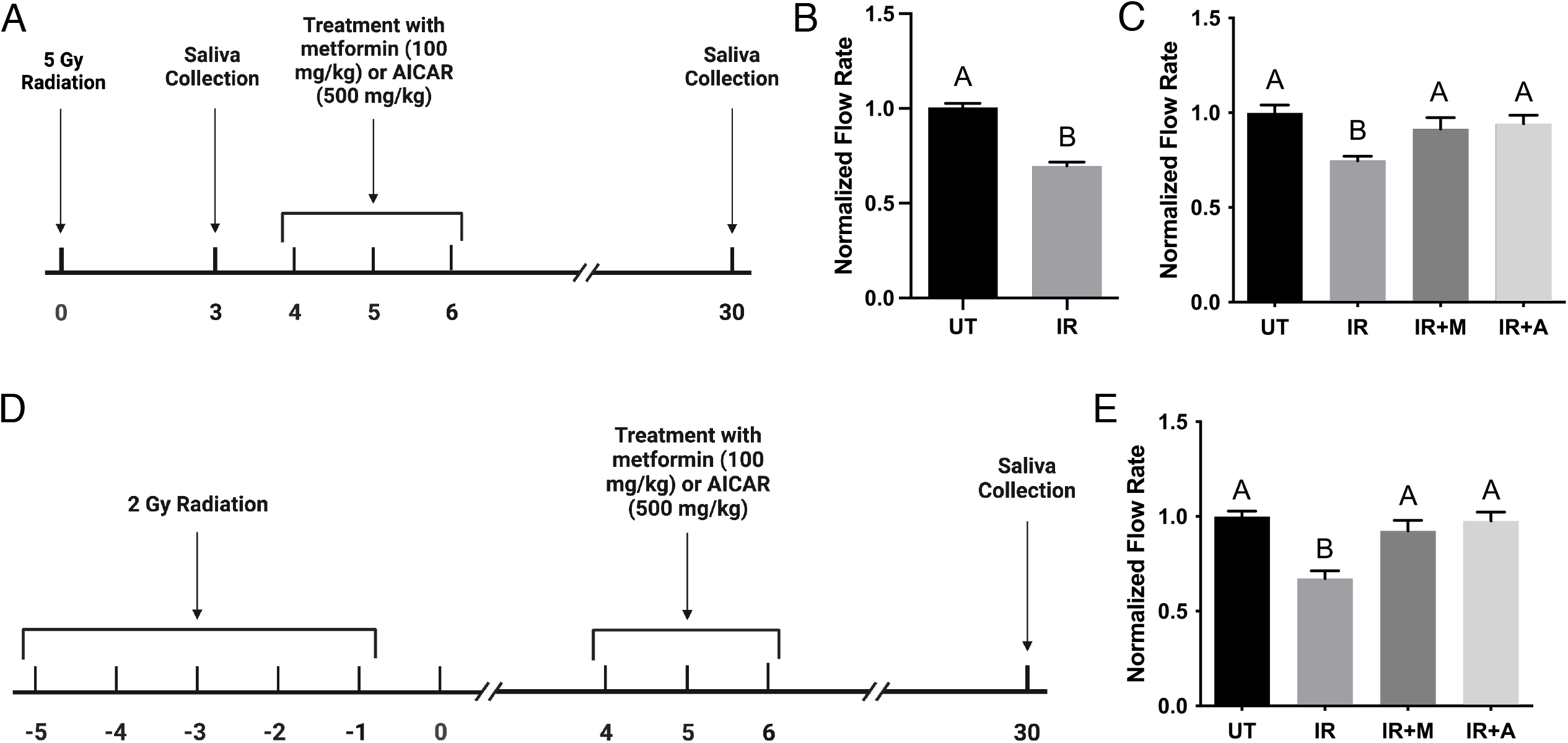
AMPK activation with AICAR or Metformin increases salivary output following radiation. (A, D) Timeline of radiation and AICAR or metformin administration for salivary output experiments following (A) single-day radiation (5-Gy x 1 day) exposure or (D) fractionated radiation (2 Gy x 5 days) exposure. (A-B) FVB mice were randomly assigned to receive no treatment (UT; n=22) or 5-Gy radiation (IR; n=30). Carbachol-stimulated saliva (0.25 mg/kg) was collected 3 days following radiation. (C) Irradiated mice were then randomized to receive three doses of either AICAR (500 mg/kg; n=10) or Metformin (100 mg/kg; n=8) on days 4, 5, and 6 following radiation, or no further treatment (n=12) and saliva was collected 30 days following radiation. (D-E) FVB mice were randomly assigned to receive no treatment (UT; n=10) or 2 Gy fractionated radiation (IR; n=30). Irradiated mice were then randomized to receive three doses of either AICAR (500 mg/kg; n=10) or Metformin (100 mg/kg; n=10) on days 4, 5, and 6 following the last dose of radiation, or no further treatment (n=10) and saliva was collected 30 days following the last dose of radiation. Data are combined from multiple independent experiments and presented as mean ± SEM. Statistical differences were determined with a one-way ANOVA with Newman-Keuls multiple comparisons test, p<0.05.

To evaluate the role of AMPK activation in the re-differentiation response of salivary glands to radiation, we evaluated the proportion of amylase producing cells 30 days following radiation treatment. Radiation treatment decreases amylase-positive area by 13% compared to UT (Fig. 5A-E). Treatment with AICAR or Metformin at days 4, 5, and 6 following radiation restores amylase area to levels observed in unirradiated mice (Fig. 5A-E). Additionally, radiation treatment decreases a marker of apical/basolateral polarity (phosphorylated aPKCζT560-positive area) by 10% compared to UT (Fig. 5F-J). Treatment with AICAR or metformin increases phosphorylated aPKCζ^T560^ area, albeit levels in metformin mice are lower than unirradiated mice (Fig. 5F-J). To further assess the effect of pharmacological AMPK activation on parotid gland proliferation, we counted Ki67+ cells 30 days post-IR with and without metformin or ACIAR treatment. Radiation increases Ki67+ parotid cells by 65% at day 30 following IR compared to UT (Fig. 5K-O), and treatment with AICAR or metformin reduces Ki67+ parotid cells to UT levels (Fig. 5K-O). Taken together, these data implicate a role for AMPK activation in restoring salivary gland function following radiation.

**Figure 5.**
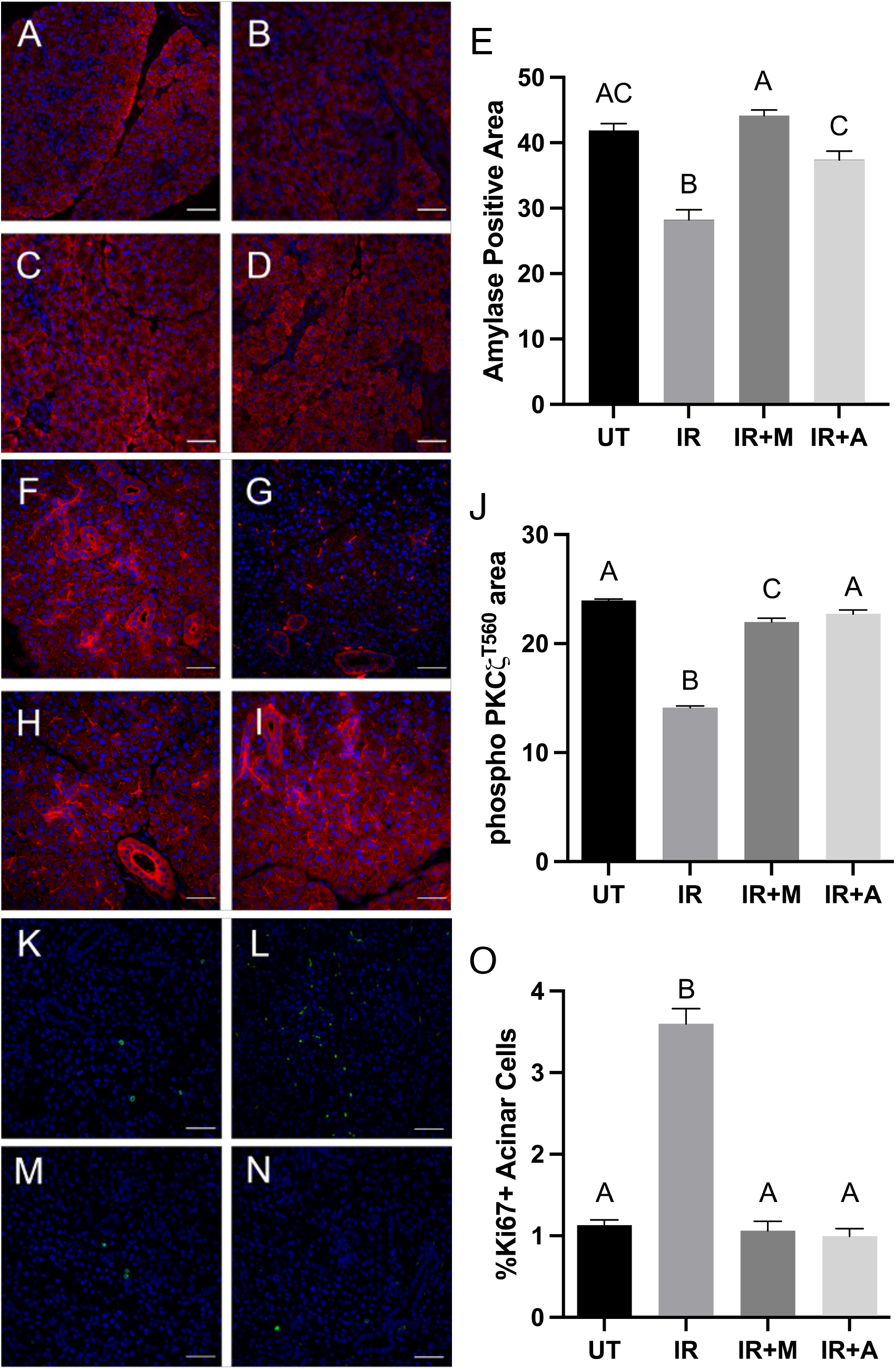
AMPK activation with AICAR or metformin increases amylase and phosphorylated aPKCζ area and decreases proliferation following radiation. FVB mice were untreated or exposed to 5-Gy radiation with a subset of mice receiving metformin (100mg/kg) or AICAR (500mg/kg) on days 4, 5 and 6 following radiation. Parotid salivary glands were collected 30 days following radiation and immunofluorescence for amylase, phosphorylated aPKCζT560, or Ki67 was performed. (A-D) Representative images of amylase positive acinar cells (red) in (A) untreated (n=4), (B) 5-Gy radiation (n=4), (C) 5-Gy radiation plus metformin (n=4), and (D) 5-Gy radiation plus AICAR (n=4). (E) The percentage of amylase positive area was determined using ImageJ software from at least 20 fields of view per mouse. The graph represents the percentage of positive amylase area as a percentage of the total area. (F-I) Representative images of phosphorylated aPKCζ positive acinar cells (red) in (F) untreated (n=3), (G) 5-Gy radiation (n=3), (H) 5-Gy radiation plus metformin (n=3), and (I) 5-Gy radiation plus AICAR (n=3). (J) The percentage of positive phosphorylated aPKCζ was determined using ImageJ software from at least 20 fields of view per mouse. The graph represents the percentage of positive phosphorylated aPKCζ as a percentage of the total area. (J) Graphs represent the mean ± SEM. Significant differences were determined via a one-way ANOVA followed by Bonferroni’s post-hoc comparisons. (K-N) Representative images of Ki67 positive acinar cells (green) over total nuclei stained with DAPI (blue) in (K) untreated (n=5), (L) 5-Gy radiation (n=5), (M) 5-Gy radiation plus metformin (n=5), and (N) 5-Gy radiation plus AICAR (n=5). (O) Quantification of Ki67+ nuclei as a percentage of the total number of nuclei from 5 fields of view per mouse. Data are presented as mean ± SEM and statistical differences were determined with a one-way ANOVA followed by Tukey’s post-hoc test, p<0.05.

## Discussion

Chronic hyposalivation remains an incurable side effect of radiation therapy; therefore, research into novel therapeutic targets is necessary. In the present study, we found that AMPK activation was decreased at days 3, 5, and 7 following IR, under conditions of altered cellular energy homeostasis evidenced by decreased intracellular NAD+ levels and AMP:ATP ratio (Fig. 1). Post-radiation treatment with pharmacological AMPK activators, AICAR or metformin, improved salivary output 30 days following radiation (Fig. 4C). This work provides a potential new model utilizing AMPK activation for functional salivary gland restoration that may prove safer and more cost-effective than some proposed therapies, including gene therapy and surgical strategies (Sasportas et al. 2013; Sood et al. 2014). Although research utilizing metformin in a salivary gland damage model hasn’t been described, metformin has been widely studied in other models of epithelial wound healing. Han et al. found significantly improved cutaneous wound healing in diabetic mice treated with metformin (Han et al. 2017). Further, in an aged-mouse model of cutaneous would healing, metformin, as well as AMPK activator resveratrol, improved the speed of cutaneous wound healing (Zhao et al. 2017). Activation of AMPK by metformin and resveratrol likely has multiple consequences, including mTOR inhibition, autophagy induction and subsequent inhibition of protein synthesis and proliferation. Persistent compensatory proliferation in irradiated salivary glands is associated with negative functional outcomes (Chibly et al. 2018; Grundmann et al. 2010; Morgan-Bathke et al. 2015); inhibition of proliferation and associated anabolic processes may improve cell homeostasis and restore acinar cell function in irradiated salivary glands.

AMPK activation leads to inhibition of mTOR activity through phosphorylation of TSC2 and Raptor and activation of ULK1 which collectively stimulates autophagy. Decreases in AMPK activation (Fig. 1) aligns with previous work demonstrating increased mTOR activity and lack of autophagy induction following IR (Morgan-Bathke et al. 2015). Autophagy-deficient mice have exacerbated radiation-induced salivary gland dysfunction and treatment with a rapalogue (CCI-779) that inhibits mTOR activity restores salivary gland function (Morgan-Bathke et al. 2014; Morgan-Bathke et al. 2015). Correspondingly, activation of AMPK and inhibition of mTOR lead to reductions in compensatory proliferation and restoration of salivary gland function (Fig 4 and Morgan-Bathke et al. 2015). Intriguingly, treatment with CCI-779 or metformin alone reduced salivary secretion at day 30. In the case of CCI-779, secretion normalized at day 60 and we postulate metformin would have the same result. Additionally, AMPK activation has been shown to regulate mitophagy and the removal of defective mitochondria (Herzig and Shaw 2018). We have recently demonstrated that radiation leads to a reduction of genes involved in lipid β-oxidation and oxidative phosphorylation suggestive of a mitochondrial dysfunction phenotype (Meeks et al. 2021). Therefore, restoration of salivary gland function by AICAR or metformin may involve the stimulation of mitophagy and subsequent mitochondrial biogenesis in order to restore mitochondrial function.

Recently the LKB1-AMPK signaling axis has been shown to be intertwined with regulation of adherens and tight junctions, apical/basolateral polarity and actin cytoskeleton assembly (Tsukita et al. 2019; Zhu et al. 2018). In cell culture models, LKB1 localization at adherens junctions has been shown to be critical in activation of AMPK localized at tight junctions (Sebbagh et al. 2009; Zhang et al. 2006). In addition, pharmacological activation of AMPK via AICAR or metformin aids in the re-establishment of intestinal epithelial barrier function in experimental models of colitis or intestinal dysfunction caused by heat stress (Chen et al. 2018; Xia et al. 2019). The mechanisms of AMPK regulation of cellular junctions and polarity are currently unclear especially in the context of salivary glands. For example, AMPK activation of the transcription factor caudal type homeobox 2 (CDX2) increased expression of junctional proteins (e.g. occludin, E-cadherin) (Sun et al. 2018; Sun et al. 2017); however, this family of transcription factors is expressed at a very low level in salivary glands (Salivary Gland Atlas). Following radiation damage to salivary glands, there is decreased apical/basolateral polarity, decreased E-cadherin/β-catenin association and cytoskeletal disruptions five days after exposure (Chibly et al. 2018; Wong et al. 2018). Interestingly, decreases in AMPK activation (Fig. 1) precede this phenotype and may suggest a role in homeostatic maintenance of these structures. Treatment with AICAR or metformin following radiation increases levels of apical/basolateral polarity (Fig. 5) suggesting a re-establishment of epithelial differentiation that is critical for the restoration of saliva secretion. This mechanistic role of AMPK activation in salivary output provides additional support for clinical studies utilizing metformin or other AMPK activators and broadens treatment options available for head and neck cancer patients undergoing radiotherapy.

In summary, radiation treatment leads to reductions in key components of the AMPK signaling pathways at acute time points (days 3-5) within salivary glands, notably phosphorylated AMPK, NAD+, AMP, Sirt1 and NAMPT levels compared to untreated glands. Activation of AMPK following IR reduces compensatory proliferation, increases differentiation markers and restores both apical/basolateral polarity and salivary flow rates to levels comparable to unirradiated controls. This work provides a novel therapeutic target for functional restoration following radiotherapy that could eventually provide relief for those affected by chronic hyposalivation.

## Acknowledgements

Author 1: contributed to conception and design, data acquisition, analysis, and interpretation, and drafting and critically revising the manuscript. Author 2: contributed to data acquisition and analysis and critically revised the manuscript. Author 3: contributed to data acquisition and analysis and critically revised the manuscript. Author 4: contributed to data acquisition and analysis and critically revised the manuscript. Author 5: contributed to conception and design, data interpretation, and drafting and critically revising the manuscript. All authors gave their final approval and agree to be accountable for all aspects of the work. This work was supported by the National Institute of Dental and Craniofacial Research [DE023534 and DE029506].

The authors declare no conflict of interest.

## References

Abreu-Blanco MT, Watts JJ, Verboon JM, Parkhurst SM. 2012. Cytoskeleton responses in wound repair. Cellular and Molecular Life Sciences. 69(15):2469–2483.

Agarwal S, Bell CM, Rothbart SB, Moran RG. 2015. Amp-activated protein kinase (ampk) control of mtorc1 is p53- and tsc2-independent in pemetrexed-treated carcinoma cells. Journal of Biological Chemistry. 290(46):27473–27486.

Brandauer J, Vienberg SG, Andersen MA, Ringholm S, Risis S, Larsen PS, Kristensen JM, Frøsig C, Leick L, Fentz J et al. 2013. Amp-activated protein kinase regulates nicotinamide phosphoribosyl transferase expression in skeletal muscle. J Physiol. 591(20):5207–5220.

Cantó C, Gerhart-Hines Z, Feige JN, Lagouge M, Noriega L, Milne JC, Elliott PJ, Puigserver P, Auwerx J. 2009. Ampk regulates energy expenditure by modulating nad+ metabolism and sirt1 activity. Nature. 458(7241):1056–1060.

Chen L, Wang J, You Q, He S, Meng Q, Gao J, Wu X, Shen Y, Sun Y, Wu X et al. 2018. Activating ampk to restore tight junction assembly in intestinal epithelium and to attenuate experimental colitis by metformin. Front Pharmacol. 9:761.

Chibly AM, Wong WY, Pier M, Cheng H, Mu Y, Chen J, Ghosh S, Limesand KH. 2018. Apkcζ-dependent repression of yap is necessary for functional restoration of irradiated salivary glands with igf-1. Scientific Reports. 8(1).

Cramer JD, Burtness B, Le QT, Ferris RL. 2019. The changing therapeutic landscape of head and neck cancer. Nature Reviews Clinical Oncology. 16(11):669–683.

Garten A, Schuster S, Penke M, Gorski T, de Giorgis T, Kiess W. 2015. Physiological and pathophysiological roles of nampt and nad metabolism. Nat Rev Endocrinol. 11(9):535–546.

Gilman KE, Camden JM, Klein RR, Zhang Q, Weisman GA, Limesand KH. 2019. P2×7 receptor deletion suppresses γ-radiation-induced hyposalivation. American Journal of Physiology-Regulatory, Integrative and Comparative Physiology. 316(5):R687–R696.

Gilman KE, Camden JM, Woods LT, Weisman GA, Limesand KH. 2021. Indomethacin treatment post-irradiation improves mouse parotid salivary gland function via modulation of prostaglandin e2 signaling. Front Bioeng Biotechnol. 9:697671.

Golla S, Golla JP, Krausz KW, Manna SK, Simillion C, Beyoğlu D, Idle JR, Gonzalez FJ. 2017. Metabolomic analysis of mice exposed to gamma radiation reveals a systemic understanding of total-body exposure. Radiat Res. 187(5):612–629.

Grundmann O, Fillinger JL, Victory KR, Burd R, Limesand KH. 2010. Restoration of radiation therapy-induced salivary gland dysfunction in mice by post therapy igf-1 administration. BMC Cancer. 10(1):417.

Grundmann O, Mitchell GC, Limesand KH. 2009. Sensitivity of salivary glands to radiation: From animal models to therapies. Journal of Dental Research. 88(10):894–903.

Han X, Tao Y, Deng Y, Yu J, Sun Y, Jiang G. 2017. Metformin accelerates wound healing in type 2 diabetic db/db mice. Molecular Medicine Reports. 16(6):8691–8698.

Hao B, Xiao Y, Song F, Long X, Huang J, Tian M, Deng S, Wu Q. 2017. Metformin-induced activation of ampk inhibits the proliferation and migration of human aortic smooth muscle cells through upregulation of p53 and ifi16. International Journal of Molecular Medicine.

Hardie DG, Carling D, Carlson M. 1998. The amp-activated/snf1 protein kinase subfamily: Metabolic sensors of the eukaryotic cell? Annual Review of Biochemistry. 67(1):821–855.

Herzig S, Shaw RJ. 2018. Ampk: Guardian of metabolism and mitochondrial homeostasis. Nature Reviews Molecular Cell Biology. 19(2):121–135.

Imamura K, Ogura T, Kishimoto A, Kaminishi M, Esumi H. 2001. Cell cycle regulation via p53 phosphorylation by a 5′-amp activated protein kinase activator, 5-aminoimidazole-4-carboxamide-1-β--ribofuranoside, in a human hepatocellular carcinoma cell line. Biochemical and Biophysical Research Communications. 287(2):562–567.

Jensen SB, Vissink A, Limesand KH, Reyland ME. 2019. Salivary gland hypofunction and xerostomia in head and neck radiation patients. JNCI Monographs. 2019(53).

Jones RG, Plas DR, Kubek S, Buzzai M, Mu J, Xu Y, Birnbaum MJ, Thompson CB. 2005. Amp-activated protein kinase induces a p53-dependent metabolic checkpoint. Molecular Cell. 18(3):283–293.

Ke R, Xu Q, Li C, Luo L, Huang D. 2018. Mechanisms of ampk in the maintenance of atp balance during energy metabolism. Cell Biology International. 42(4):384–392.

Kim JW, Kim SM, Park JS, Hwang SH, Choi J, Jung KA, Ryu JG, Lee SY, Kwok SK, Cho ML et al. 2019. Metformin improves salivary gland inflammation and hypofunction in murine sjögren’s syndrome. Arthritis Res Ther. 21(1):136.

Lee IH. 2019. Mechanisms and disease implications of sirtuin-mediated autophagic regulation. Exp Mol Med. 51(9):1–11.

Limesand KH, Avila JL, Victory K, Chang HH, Shin YJ, Grundmann O, Klein RR. 2010. Insulin-like growth factor-1 preserves salivary gland function after fractionated radiation. Int J Radiat Oncol Biol Phys. 78(2):579–586.

Limesand KH, Said S, Anderson SM. 2009. Suppression of radiation-induced salivary gland dysfunction by igf-1. PLoS ONE. 4(3):e4663.

Manna SK, Krausz KW, Bonzo JA, Idle JR, Gonzalez FJ. 2013. Metabolomics reveals aging-associated attenuation of noninvasive radiation biomarkers in mice: Potential role of polyamine catabolism and incoherent DNA damage-repair. J Proteome Res. 12(5):2269–2281.

Meeks L, De Oliveira Pessoa D, Martinez JA, Limesand KH, Padi M. 2021. Integration of metabolomics and transcriptomics reveals convergent pathways driving radiation-induced salivary gland dysfunction. Physiol Genomics. 53(3):85–98.

Morgan-Bathke M, Harris ZI, Arnett DG, Klein RR, Burd R, Ann DK, Limesand KH. 2014. The rapalogue, cci-779, improves salivary gland function following radiation. PLoS ONE. 9(12):e113183.

Morgan-Bathke M, Hill GA, Harris ZI, Lin HH, Chibly AM, Klein RR, Burd R, Ann DK, Limesand KH. 2015. Autophagy correlates with maintenance of salivary gland function following radiation. Scientific Reports. 4(1).

Niessen MT, Scott J, Zielinski JG, Vorhagen S, Sotiropoulou PA, Blanpain C, Leitges M, Niessen CM. 2013. Apkcλ controls epidermal homeostasis and stem cell fate through regulation of division orientation. Journal of Cell Biology. 202(6):887–900.

Sasportas LS, Hosford AT, Sodini MA, Waters DJ, Zambricki EA, Barral JK, Graves EE, Brinton TJ, Yock PG, L. Q-T et al. 2013. Cost-effectiveness landscape analysis of treatments addressing xerostomia in patients receiving head and neck radiation therapy. Oral Surgery, Oral Medicine, Oral Pathology and Oral Radiology. 116(1):e37–e51.

Sebbagh M, Santoni M-J, Hall B, Borg J-P, Schwartz MA. 2009. Regulation of lkb1/strad localization and function by e-cadherin. Current Biology. 19(1):37–42.

Siegel RL, Miller KD, Fuchs HE, Jemal A. 2022. Cancer statistics, 2022. CA Cancer J Clin. 72(1):7–33.

Sood AJ, Fox NF, O’Connell BP, Lovelace TL, Nguyen SA, Sharma AK, Hornig JD, Day TA. 2014. Salivary gland transfer to prevent radiation-induced xerostomia: A systematic review and meta-analysis. Oral Oncology. 50(2):77–83.

Sun X, Du M, Navarre DA, Zhu M-J. 2018. Purple potato extract promotes intestinal epithelial differentiation and barrier function by activating amp-activated protein kinase. Molecular Nutrition & Food Research. 62(4):1700536.

Sun X, Yang Q, Rogers CJ, D. M, Zhu M-J. 2017. Ampk improves gut epithelial differentiation and barrier function via regulating cdx2 expression. Cell Death & Differentiation. 24(5):819–831.

Tsukita K, Yano T, Tamura A, Tsukita S. 2019. Reciprocal association between the apical junctional complex and ampk: A promising therapeutic target for epithelial/endothelial barrier function? International Journal of Molecular Sciences. 20(23):6012.

Wong WY, Pier M, Limesand KH. 2018. Persistent disruption of lateral junctional complexes and actin cytoskeleton in parotid salivary glands following radiation treatment. Am J Physiol Regul Integr Comp Physiol. 315(4):R656–r667.

Xia Z, Huang L, Yin P, Liu F, Liu Y, Zhang Z, Lin J, Zou W, Li C. 2019. L-arginine alleviates heat stress-induced intestinal epithelial barrier damage by promoting expression of tight junction proteins via the ampk pathway. Molecular Biology Reports. 46(6):6435–6451.

Zhang L, Li J, Young LH, Caplan MJ. 2006. Amp-activated protein kinase regulates the assembly of epithelial tight junctions. Proceedings of the National Academy of Sciences. 103(46):17272–17277.

Zhao P, Sui B-D, Liu N, Lv Y-J, Zheng C-X, Lu Y-B, Huang W-T, Zhou C-H, Chen J, Pang D-L et al. 2017. Anti-aging pharmacology in cutaneous wound healing: Effects of metformin, resveratrol, and rapamycin by local application. Aging Cell. 16(5):1083–1093.

Zhu M-J, Sun X, Du M. 2018. Ampk in regulation of apical junctions and barrier function of intestinal epithelium. Tissue Barriers. 6(2):1–13.

