## Appendix for "AMPK activation restores salivary function following radiation treatment"

Mice. Mice were housed in cages exposed to 12-hour light/dark cycles with *ad libitum* access to food and water. Size- and age-matched 4-8-week-old female FVB mice were randomly assigned to treatment groups for all experiments.

ATP cell assay. Primary parotid gland cells were prepared as previously described (Gilman et al. 2019). On day 3 of culture, cells were left untreated or received 5Gy radiation and were collected 72 hours later. A portion of the sample was collected for protein extraction and the remaining sample was collected for an ATP bioluminescence assay (Roche, Tucson, AZ). Intracellular ATP was extracted by adding 9 volumes of boiling 100mM Tris, 4mM EDTA (pH, 7.75) and boiling at 100°C for 2 additional minutes. ATP concentration was determined following the manufacturers’ instructions and was normalized to the protein concentration of each sample.

Western blot antibodies: anti-p-AMPK (Cell Signaling #2531, Danvers, MA, 1:500 in TBST with 5% BSA); anti-AMPK (Cell Signaling #2603, 1:500 in TBST with 5% BSA); anti-ERK1/2 (Cell Signaling #9102, 1:1000 in TBST).

RNA isolation and qRT-PCR. Parotid glands were removed from mice at days 3, 5 and 30 after 5 Gy radiation and RNA isolated as previously described (Gilman et al. 2019). Primers to NAMPT, Sirt1, and 15S ribosomal RNA were purchased from Integrated DNA Technologies (Coralville, IA). Target genes were normalized to 15S ribosomal RNA. Primer sequences: NAMPT (F: 5’-GGCTACGTGGACGACGACAC-3’; R: 5’-CATCCCCTGCAGGCCTGGTCT-3’), Sirt1 (F: 5’- AGAGTTGCCACCAACACCTCTT-3’; R: 5’- TTAGGCCAGCATTTTCTCACTGT-3’), 15S ribosomal RNA (F: 5’-ACTATTCTGCCCGAGATGGTG-3’, R: 5’-TGCTTTACGGGCTTG TAGGTG- 3’).

Histology. Salivary glands were removed, fixed in 10% (v/v) formalin for 24 hours and sent to IDEXX Bioresearch (Columbia, MO), where they were transferred to 70% (v/v) ethanol, embedded in paraffin, sectioned into 4 μm sections, and returned for immunofluorescent staining.

Immunofluorescent staining and quantification. Slides were stained with antibodies to anti-Ki67 (Cell Signaling, #9129S, Danvers, MA, diluted 1:400 in 1% BSA in PBS), anti-amylase (Sigma Aldrich, diluted 1:1000 in 1% BSA in PBS), or anti-phospho-aPKCζ^T560^ (Abcam, Cambridge, United Kingdom, diluted 1:250 in 1% BSA in PBS) as previously described (Chibly et al. 2018; Gilman et al. 2021). At least 5 images per mouse (4-5 mice/group) were manually counted for Ki67-positive cells vs. the total number of cells. At least twenty fields of view per mouse (3-4 mice/group) were analyzed using ImageJ software for amylase and phosphorylated aPKCζ^T560^ positive area.

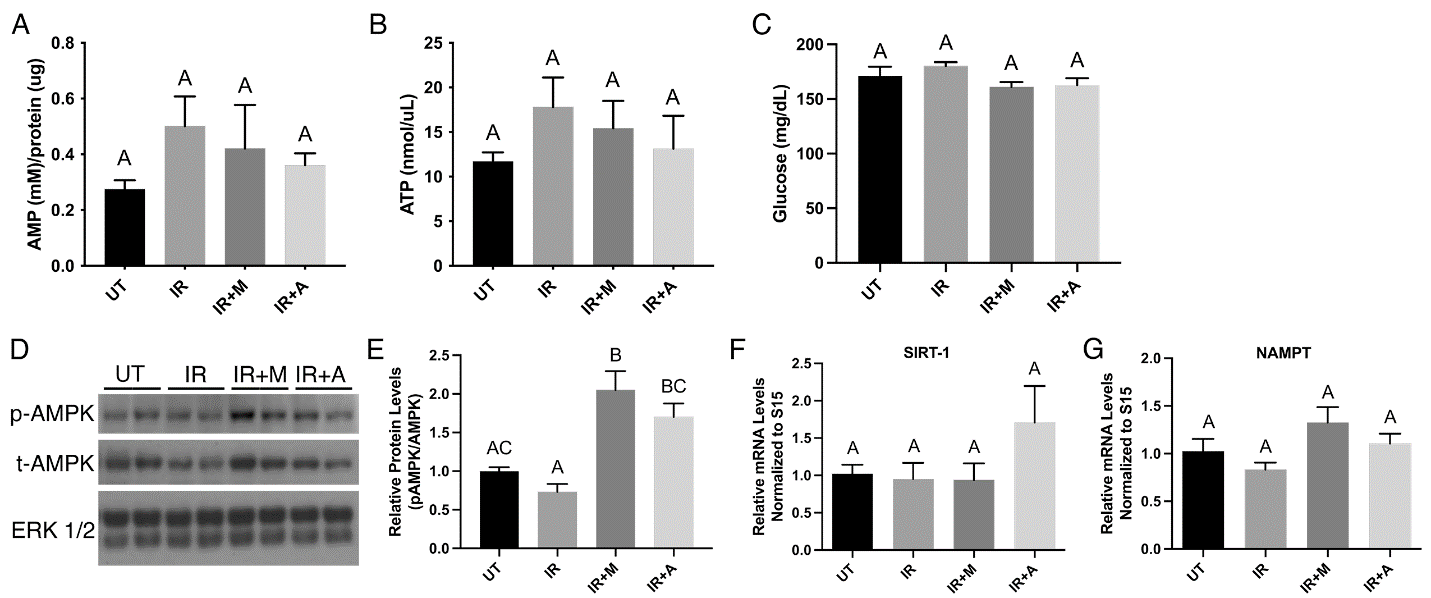

**Supplemental Figure 1. AICAR or metformin treatment increase AMPK phosphorylation at day 30 following IR and have no effect on blood glucose, AMP and ATP levels, or expression of SIRT1 and NAMPT.** FVB mice were untreated or exposed to 5-Gy IR, with select mice receiving Metformin or AICAR injections on days 4, 5, and 6. (A, B, D-G) Parotid glands were harvested from indicated treatment groups at day 30 and levels of (A) AMP and (B) ATP were evaluated from tissue lysates (n=4/group) via colorimetric assays as described in Materials and Methods. (C) Blood was collected from indicated groups at day 30 following IR and glucose was measured following a 4-hour fast with a handheld glucometer. (D-E) Protein was extracted from parotid glands of each indicated treatment (n=4/group) and immunoblots were performed. Levels of phosphorylated AMPK (p-AMPK), total AMPK (t-AMPK) and ERK 1/2 (loading control) were evaluated via immunoblot. AMPK (p-AMPK) levels were normalized to total AMPK (t-AMPK) levels. (F-G) qRT-PCR was performed with primers specific to (F) Sirt-1 and (G) NAMPT, with data normalized to 15S ribosomal RNA as an internal control and calculated as fold-change relative to the average of untreated mice (n=4/group). (A-G) Data are presented as mean ± SEM. Statistical differences were determined with a one-way ANOVA followed by Tukey’s post-hoc test. Groups with different letter designations are significantly different from each other, p<0.05.

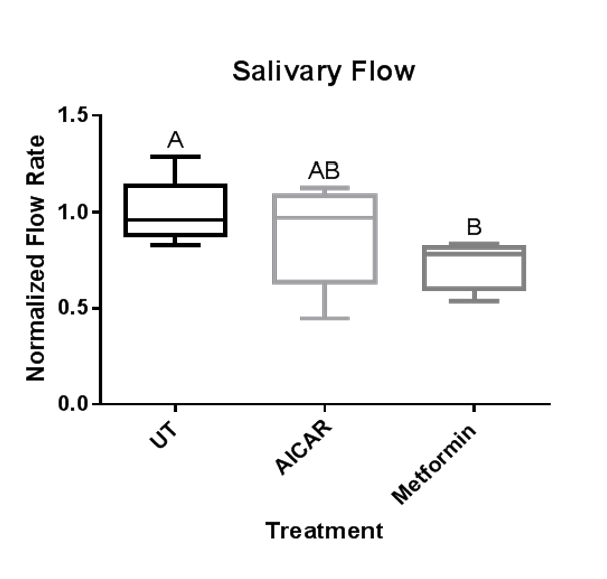

**Supplemental Figure 2. Metformin decreases salivary output in non-irradiated mice.** FVB mice were randomly assigned to receive no treatment (UT, n=18), three injections of AICAR (500 mg/kg, n=8), or three doses of metformin (100 mg/kg; n=8). Carbachol-stimulated saliva (0.25 mg/kg) was collected 24 days following the final dose. Data are presented as means ± SEM and statistical differences were determined with one-way ANOVA with Newman-Keuls multiple comparisons test, p<0.05.
