## Supplementary figures and images for "AMPK activation restores salivary function following radiation treatment"

### Supplemental Figure 1

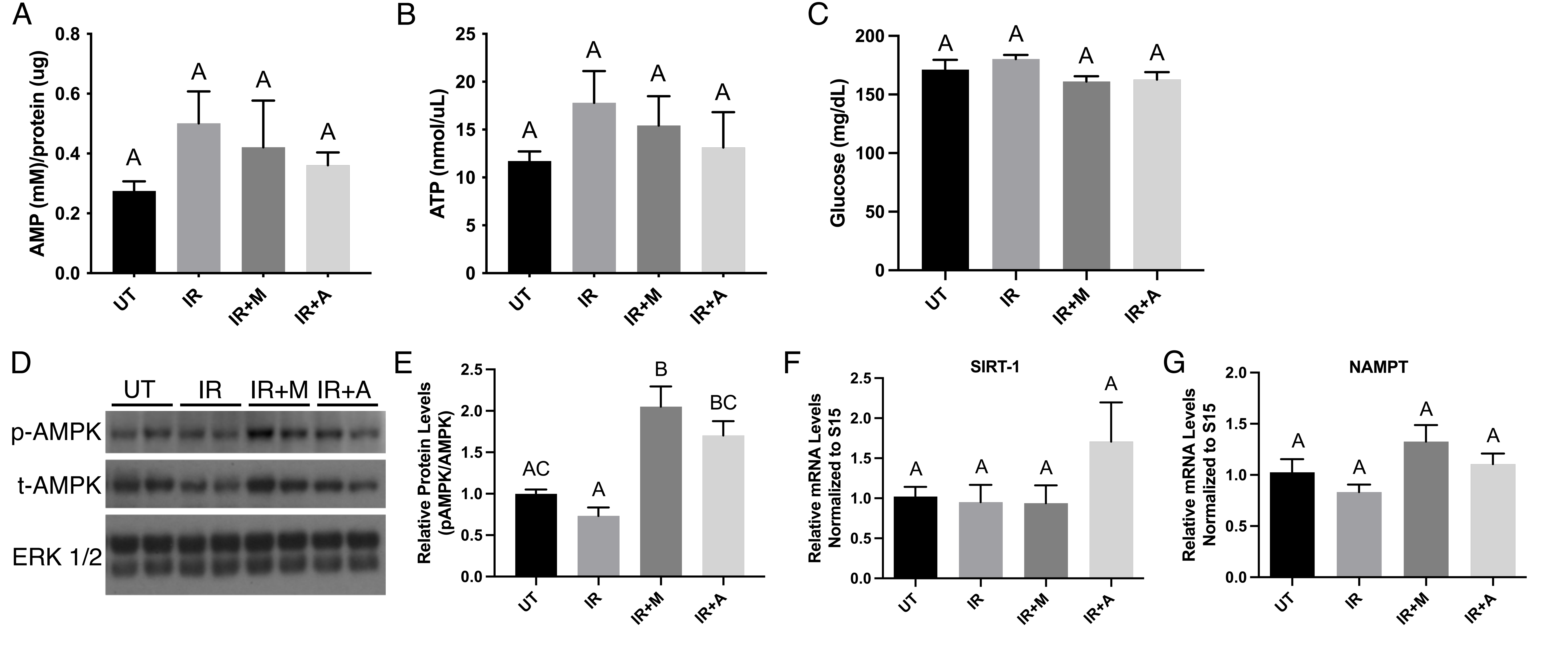

### Supplemental Figure 2

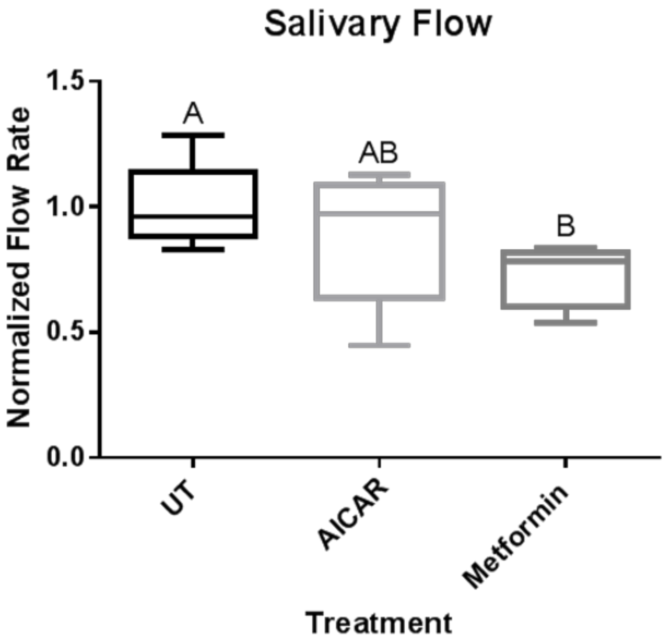
